# Effects of seasonal changes on the carbon dynamics in mixed coniferous forests

**DOI:** 10.1101/2022.04.08.487629

**Authors:** Tong Gao, Xinyu Song, Yunze Ren, Hui Liu, Xibin Dong

## Abstract

We characterized seasonal changes in the residual rate and mass-loss rate of litter, as well as the carbon release dynamics of litter and soil, in mixed coniferous forests in the Xiaoxinganling region by conducting a controlled freeze–thaw experiment. The carbon release rate and mass-loss rate of litter at two levels of decomposition (undecomposed and semi-decomposed) were measured during four seasons: the unfrozen season, freeze–thaw season, frozen season, and thaw season. The temperature was higher, the mass-loss rate was faster, and the overall mass-loss rate of litter was higher in the unfrozen season than in the other three seasons; litter organic carbon increased and soil organic carbon increased due to the strong carbon sequestration capacity of plants. The temperature fluctuated above and below 0°C during the freeze–thaw season, which results in the physical breakage of the undecomposed litter and increases in the mass-loss rate. This leads to increases in the organic carbon of undecomposed litter and decreases in the soil organic carbon of undecomposed litter; the opposite patterns were observed for changes in the organic carbon of semi-decomposed litter and soil organic carbon. There was noticeable mass loss of litter during the frozen season, and the rate of mass loss of litter during the thaw season was the lowest. Litter organic carbon decreased and soil organic carbon increased in both seasons. The organic carbon of undecomposed litter was highest in the thaw season, followed by the freeze–thaw season, frozen season, and unfrozen season. The organic carbon of semi-decomposed litter was highest in the frozen season, followed by the thaw season, freeze–thaw season, unfrozen season to freeze–thaw season, frozen season, thaw season, and unfrozen season. After freeze–thaw treatment, the organic carbon in deadfall soil was highest in the unfrozen season, freeze–thaw season, frozen season, thaw season to unfrozen season, frozen season, thaw season, and freeze–thaw season. The findings of this study provide new insights into the material cycling process under freeze–thawing, as well as information on the effect of seasonal freeze–thaws on the forest carbon cycle.

## Introduction

Seasonal freeze–thaw cycles affect the majority of the Earth’s mid-latitudes [1]; at mid-to-high latitudes and high altitudes, these cycles have a substantial effect on forest ecosystem processes [2,3]. Seasonal freeze–thaw cycles can last up to 5–6 months per year in Northeast China, which is one of China’s primary freeze–thaw erosion areas. The freeze–thaw cycle is particularly active in Heilongjiang Province, which is located in China’s alpine region [4]. Forests play a key role in regulating the global biosphere’s carbon balance. On the one hand, forests can absorb and fix large amounts of carbon from the atmosphere, which makes them important sinks of atmospheric CO2. On the other hand, the harvesting of forests can release fixed carbon, which makes them an important source of atmospheric CO2 [5,6]. Forest ecosystems, which are the backbone of the terrestrial biosphere, sustain a substantial carbon pool, including approximately 86% and 73% of the world’s carbon contained in vegetation and soil, respectively [7]. Forests comprise approximately 26% of the world’s surface area yet hold over 80% of the carbon stores of terrestrial vegetation, and forest ecosystems fix roughly two-thirds of the carbon in terrestrial ecosystems annually [8]. Forest litter is the principal nutrient reservoir in forest ecosystems and is produced by the metabolism of forest plants during their growth and development [9]. Forest litter is an important aspect of the material cycle of forest ecosystems and thus an important component of forest ecosystems [10]. The rate of apoplankton decomposition and conversion reflects the rate of nutrient return and nutrient cycling in forests [11]. Physical damage, microbial activity, waterlogging, and their combined effect throughout the freeze–thaw season are the main causes of net mass loss and nutrient release during the decomposition process [12]. The decomposition and conversion of apoplankton are some of the most significant biological processes in the material cycle of forest ecosystems and are critical for the creation of soil organic matter and nutrient release [13]. Forest ecosystems sustain important ecological processes in some temperate areas and at high latitudes. Soil freezing and thawing are common in temperate, high-latitude, and high-altitude ecosystems during late winter and early spring [14]. The frequency of soil freezes has reduced in some temperate areas because of global warming [15], but these changes in temperature and moisture have a substantial effect on soil organic carbon. Warming affects soil organic carbon through its effect on plant growth, which alters the number of plant residues returned to the soil, the pace of organic carbon decomposition, and the amount of organic carbon released from the soil [16]. Many studies have shown that climate change can affect litter decomposition [17,18], the amount of organic carbon released from litter [19,20], and the carbon cycle of entire ecosystems [21–23]. Litter has a major effect on the organic carbon in soil [24,25].

Here, we used a controlled freeze–thaw environment to investigate the residual rate and mass-loss rate of dead litter, as well as the response of organic carbon and soil organic carbon to seasonal changes in the undecomposed layer and semi-decomposed layer during four seasons, the unfrozen season, freeze–thaw season, frozen season, and thaw season, to evaluate the effect of seasonal freeze–thawing on the forest carbon cycle.

## Materials and Methods

### Overview of the study area

The experiment was conducted at Dongfanghong Forestry Field, Dailing Forestry Bureau, Xiaoxinganling Region, Heilongjiang Province, China, which is located 13.9 km southeast of Dailing District (46°50’8”-47°21’32”N, 128°37’46”-129°17’50”E). The site has an average slope of 10°, an average altitude of 600 m, and annual precipitation of 660 mm. The average annual temperature is 2.7°C. Summers are humid, cool, and wet, and winters are dry, cold, and snowy.

### Experimental procedures

Three 30 m × 30 m plots were established, and three 1 m × 1 m litter samples were collected at random locations within each plot; litter and soil samples were collected separately. The litter was collected from two layers: the undecomposed layer (fresh litter showing slight color change, an intact structure, and no symptoms of decomposition) and the semi-decomposed layer (darkened litter, damaged structure, and most litter decomposed). The soil surface was cleansed with dead material before sampling, and samples (length and width of 20 cm) were taken from the 0–10 cm layer.

Incubation boxes with 50.00 g of soil were established in the laboratory. In each sample plot, three replicate incubation boxes were built for each temperature cycle, and deadfall was placed on the soil surface for incubation. The amount of deadfall added to the three replicates in the incubator was 2.00 g, 3.00 g, and 4.00 g to minimize the effect of deadfall quality on organic carbon. The actual soil moisture content measured at each time point was used to calculate the soil moisture content. Temperature disparities were set for temperature cycling based on previously collected temperature data.

The experiment was conducted over four seasons, and the average temperature of the four periods was used as the temperature for the experiment based on previous temperature monitoring. Minimum and maximum temperatures of the unfrozen season (July), freeze–thaw season (October), frozen season (January), and thaw season (March) were 17°C and 30°C, −5°C and 5°C, −30°C and −16°C, and 5°C and −5°C, respectively. Incubation lasted for 12 h with 30 cycles in each season, and the boxes were withdrawn 0, 1, 3, 5, 7, 10, and 30 times in each season. Separate cassettes were set up for each temperature cycle, totaling 168 cassettes for the four seasons, to prevent the samples in the cassettes from being damaged during sampling (4 seasons × 2 deadfall species × 7 freeze–thaws times × 3 replicates). The culture boxes were removed after the number of temperature cycles had been completed; the soil in the boxes was then air-dried and crushed through a 0.15-mm sieve, and the dead material was completely removed from the culture boxes. The samples were cleaned of surface soil particles and impurities, dried to constant weight, and weighed; they were then ground and passed through a 0.15-mm sieve prior to analysis.

**Table 1:** Content of dead litter and initial soil fractions at different levels of decomposition during different seasons.

| Season | Degree of decomposition of deadfall | Organic carbon from deadfall<br>g/kg | Soil organic carbon<br>g/kg |
| --- | --- | --- | --- |
| Unfrozen season | Undecomposed | 358.4±15.29 | 52.7±12.04 |
|  | Semi-decomposed | 370.5±20.55 | 52.7±12.04 |
| Freeze–thaw season | Undecomposed | 490.9±18.82 | 45.1±5.5 |
|  | Semi-decomposed | 483.3±11.47 | 45.1±5.5 |
| Frozen season | Undecomposed | 462.5±12.08 | 42.4±4.83 |
|  | Semi-decomposed | 512.6±8.85 | 42.4±4.83 |
| Thaw season | Undecomposed | 513.5±7.9 | 39.7±6.66 |
|  | Semi-decomposed | 503.9±8.45 | 39.7±6.66 |

Note: Data in the table are mean ± standard deviation, n=3.

### Data calculations

Residual rate of litter mass:

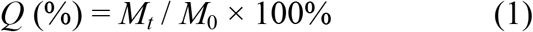

Mass loss of deadfall at all stages:

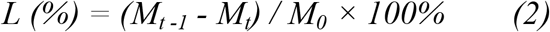

Rate of deadfall mass loss by stage (in counts):

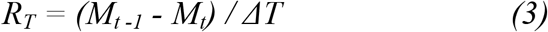

where *M*_0_ is the initial mass of the litter; *M_t_* is the residual amount of litter at each sampling event; (*M*_*t*-1_ – *M_t_*) is the difference in the residual amount of litter between two adjacent litter sampling events after drying and weighing (note: *M*_*t*-1_ is Mt measured in the previous period); and *ΔT* is the number of intervals between two adjacent sampling events.

### Analytical statistics

We used potassium dichromate-concentrated sulphuric acid with thermal oxidation to determine the amount of litter and soil organic carbon. Data analysis was conducted using IBM SPSS Statistics 26, and graphs were built using Origin 2019b. The effects of season, degree of litter decomposition, and the number of freeze–thaw cycles on the residual amount of litter, litter, and soil organic carbon were determined using repeated-measures ANOVA.

## Results

### Effect of seasonal changes on deadfall quality loss

The number of freeze–thaws had a significant effect (P<0.05) on the mass loss of deadfall (Table 2). Fig 1 shows that the rate of decrease in undecomposed litter mass residues from initial values was highest in the unfrozen season (15.89%), followed by the freeze–thaw season (13.33%), frozen season (10.56%), and thaw season (7.78%); the rate of decrease in semi-decomposed litter mass residues from initial values was highest in the unfrozen season (20.33%), frozen season (10.78%), freeze–thaw season (7.22%), and thaw season (7.22%). In the unfrozen season, the mass residual rate of undecomposed deadfall under the 3rd and 5th freeze–thaw treatments was significantly lower than the initial value; the mass residual rate of semi-decomposed deadfall under the 1st, 3rd, 5th, and 7th freeze–thaw was significantly lower than the initial value; and the mass residual rate under the 15th freeze–thaw was significantly lower than that under the 1st and 5th freeze–thaw treatments. The mass residual rate of undecomposed deadfall was significantly lower than its initial value after the 3rd and 15th freeze–thaw treatments in the freeze–thaw season, and the number of freeze–thaw treatments of semi-decomposed deadfall had no significant effect on the mass residual rate. The mass residual rate of undecomposed deadfall in the 5th and 15th freeze–thaw treatments and semi-decomposed deadfall in the 3rd, 5th, and 15th freeze–thaw treatments was much lower than initial values during the frozen season. The number of freeze–thaws that occurred throughout the thaw season had no effect on the bulk residue rate. In the unfrozen and frozen seasons, the rate of decrease in the quality of undecomposed litter was lower than that of semi-decomposed litter, and the rate of decrease in the quality of undecomposed litter was higher than that of semi-decomposed litter in the freeze–thaw season and thaw season.

**Fig 1.**
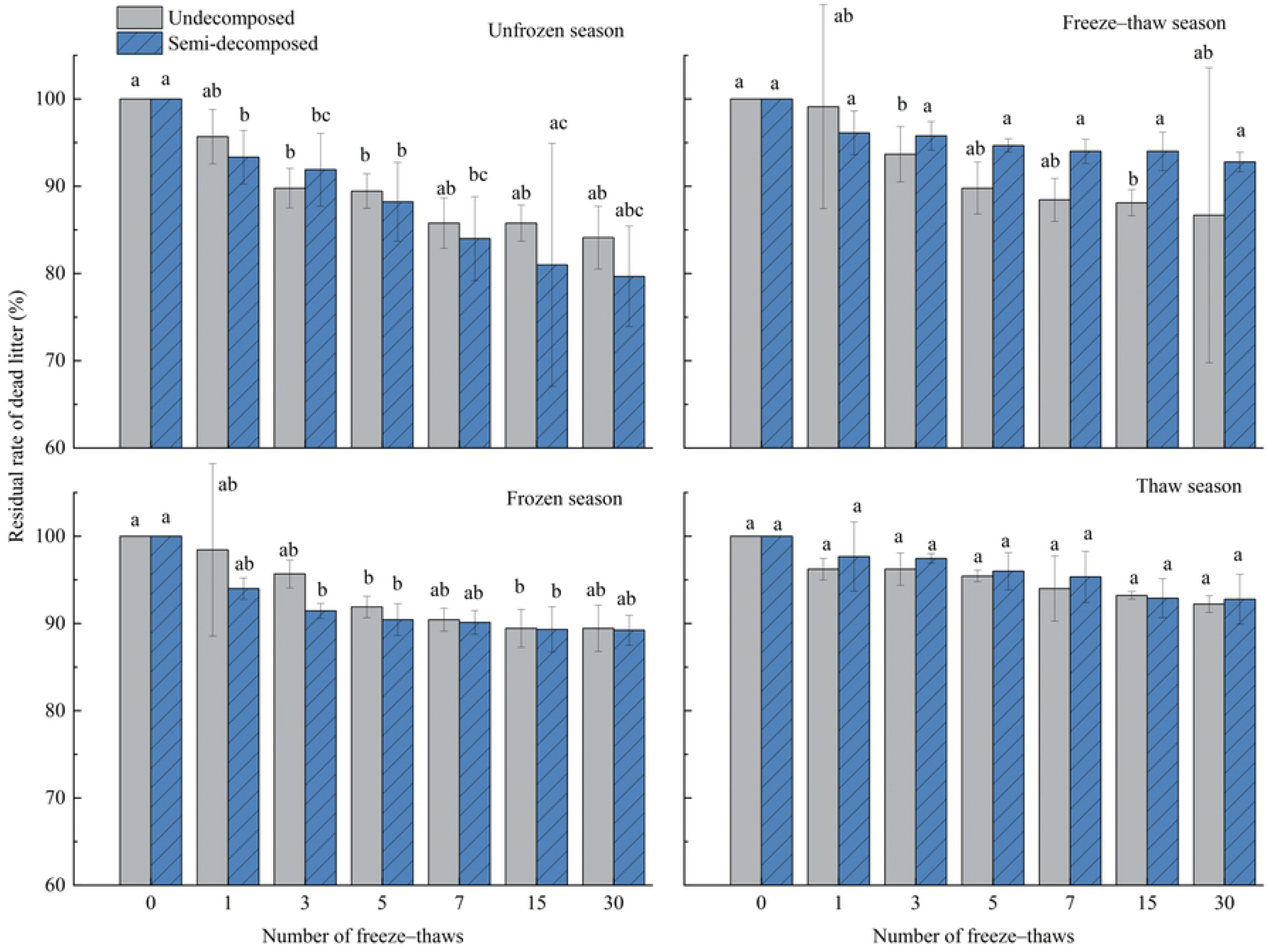
Effects of freeze–thaws in various seasons on the residual rate of litter at various levels of decomposition. Significant differences in the number of freeze–thaws (*P*<0.05) are indicated by different lowercase letters; data in the graph are means with standard deviation (n=3).

**Table 2.**
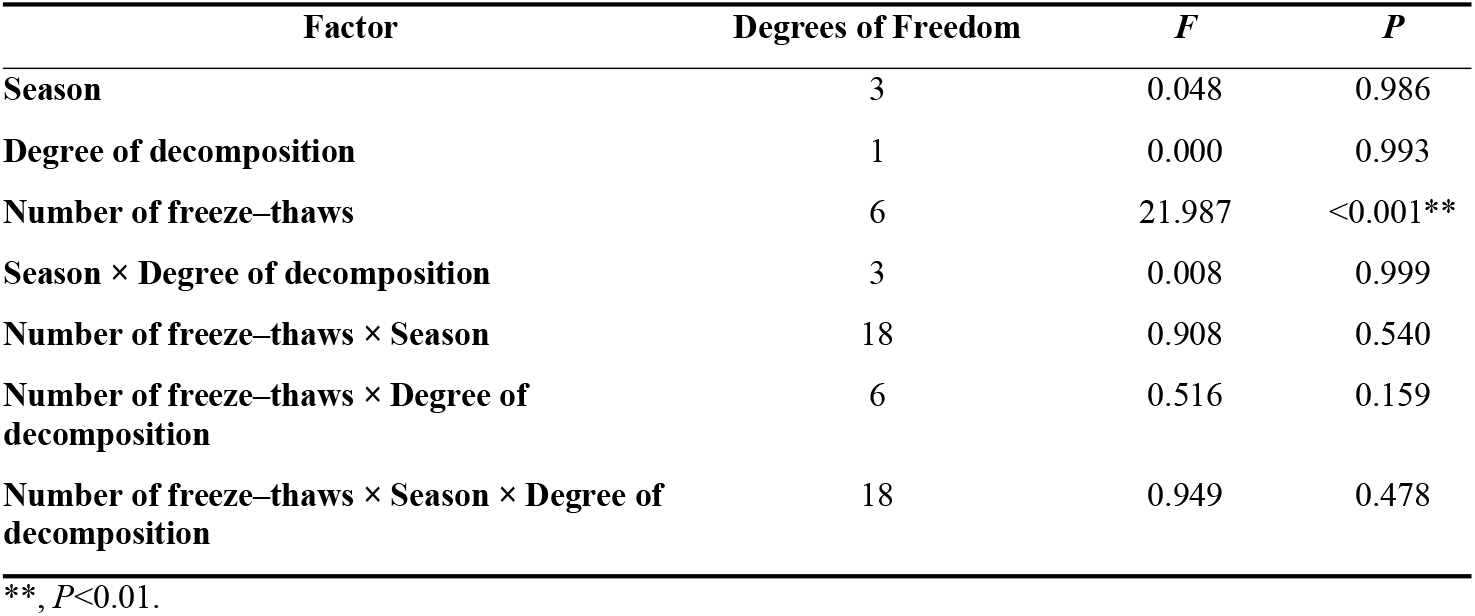
Results of repeated-measures ANOVA of litter mass loss at various degrees of decomposition as a function of various factors.

Fig 2 shows that the overall rate of deadfall mass loss was higher in the unfrozen season than in the other three seasons. The rate of mass loss of deadfall first decreased, increased, and then decreased as the number of freeze–thaw cycles increased in the freeze–thaw season and thaw season. The rate of mass loss of undecomposed deadfall in the freeze–thaw seasons first increased and then decreased, and the rate of mass loss of semi-decomposed deadfall first decreased, increased, and then decreased; the rate of mass loss of semi-decomposed deadfall was much higher after the first freeze–thaw than after other freeze–thaw intervals. During the frozen season, the rate of mass loss of undecomposed litter increased and then decreased, and the rate of semi-decomposed litter gradually decreased.

**Fig 2.**
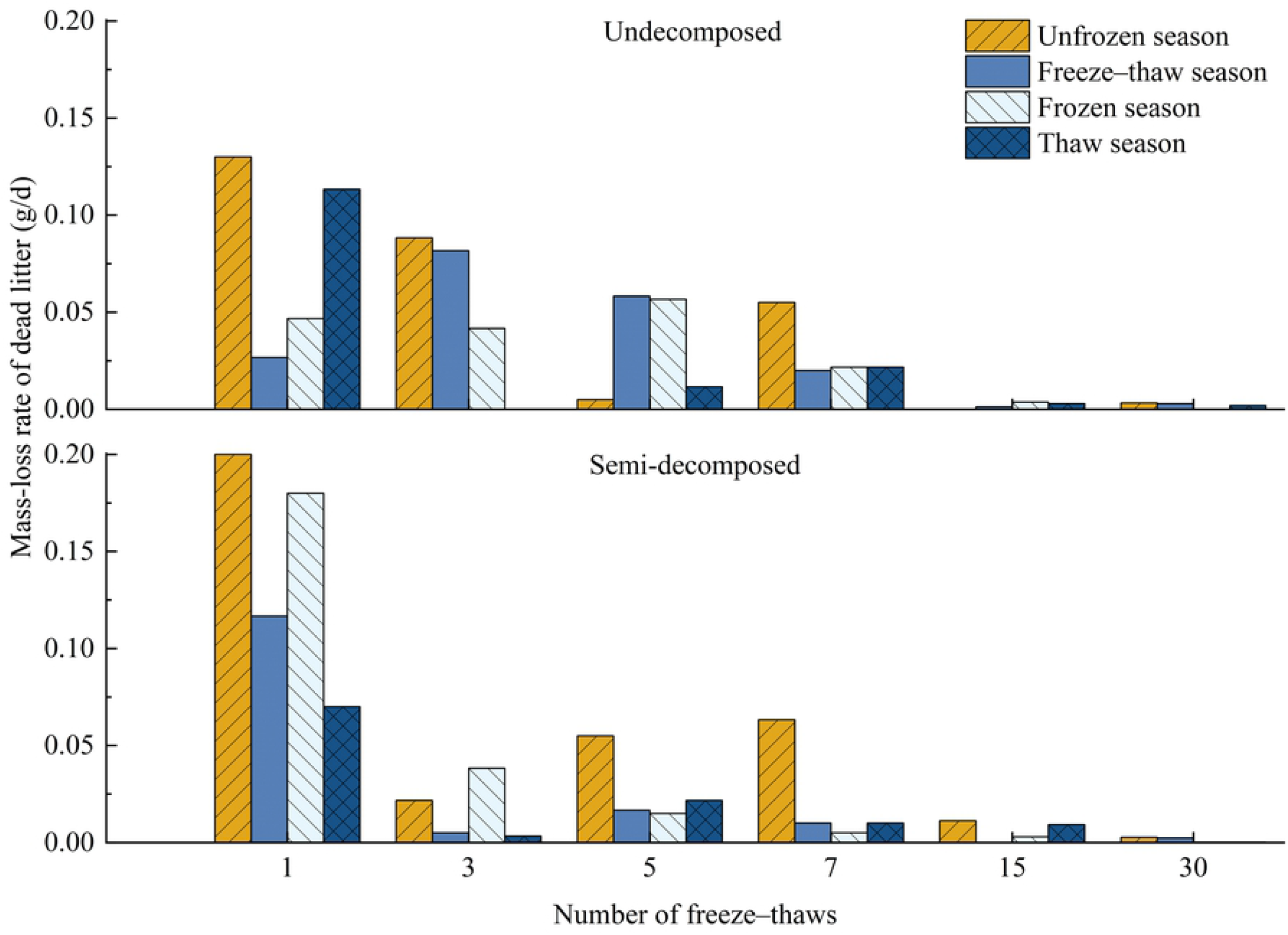
Effect of season on litter mass-loss rates at various levels of decomposition under freeze–thaw treatment.

### Effect of seasonal changes on organic carbon dynamics in litter

Table 3 shows that the number of freeze–thaws and season had a significant effect (P<0. 05) on the organic carbon of litter. Figure 3 shows that the organic carbon content of deadfall increased in the unfrozen season and decreased in the frozen and thaw seasons. The organic carbon of undecomposed deadfall was lower than the initial value in the freeze–thaw season, and the opposite was the case for semi-decomposed deadfall after 30 freeze–thaws. The interaction between the decomposition degree of deadfall and season had a significant effect on deadfall organic carbon (P=0.001). Changes in the organic carbon of litter at different decomposition levels varied after 30 freeze–thaw cycles. The organic carbon of undecomposed and semi-decomposed litter increased by 18.15% and 11.27% during the unfrozen season, respectively, and the organic carbon of undecomposed and semi-decomposed litter decreased and increased by 2.53% and 3.56% during the freeze–thaw season, respectively. The organic carbon of undecomposed litter and semi-decomposed litter decreased by 3.89% and 4.25%, respectively, during the frozen season. The organic carbon of undecomposed litter and semi-decomposed litter decreased by 5.93% and 7.46%, respectively, during the thaw season. During the frozen season, the organic carbon of semi-decomposed litter was substantially higher than that of undecomposed litter; specifically, the organic carbon of undecomposed litter was 10.12% lower than that of semi-decomposed litter.

**Fig 3.**
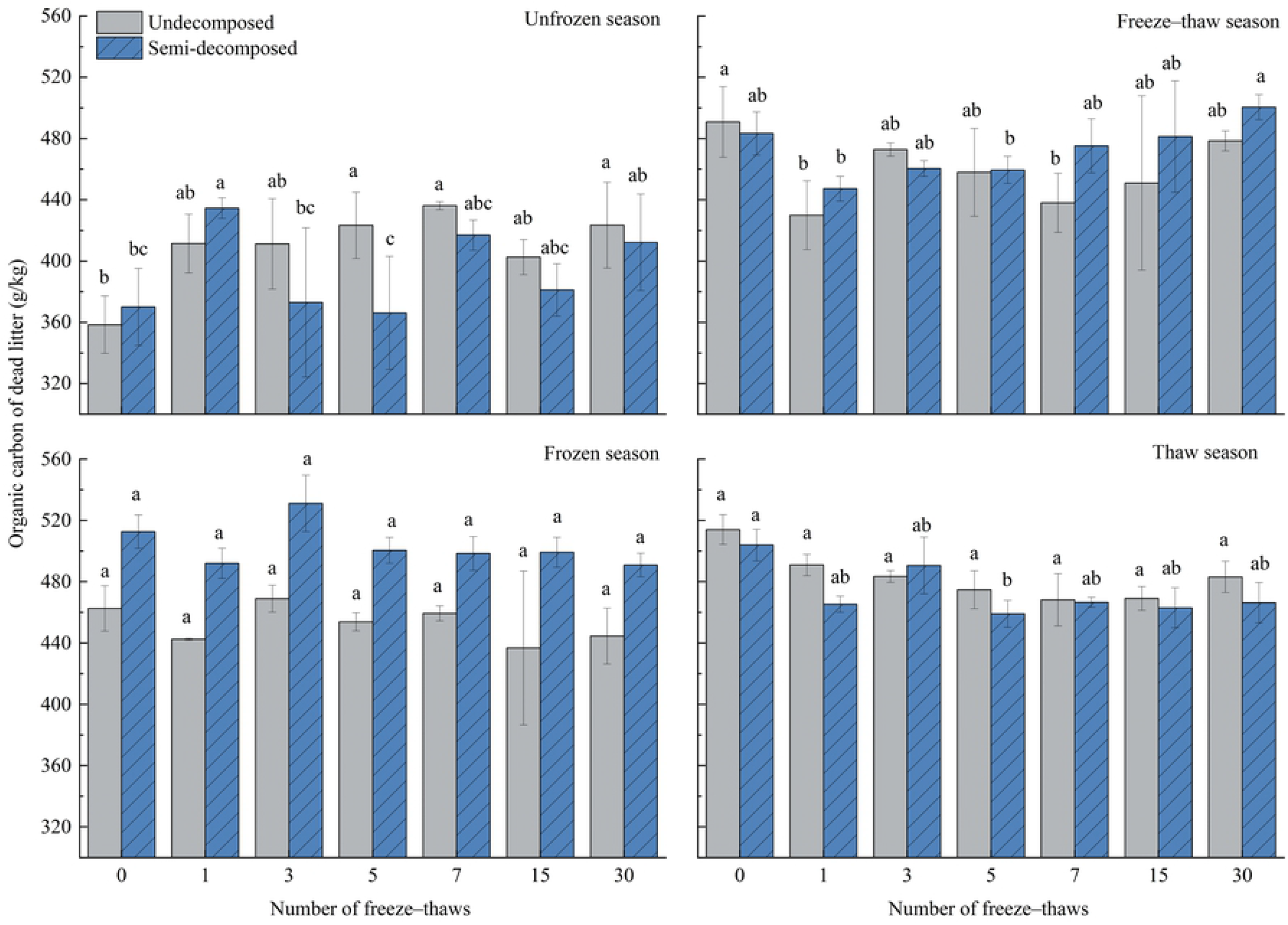
Effects of freeze–thaw time in various seasons on the carbon content of litter at various levels of decomposition. Significant differences in the number of freeze–thaws (*P*<0. 05) are indicated by different lowercase letters; data in the graphs are means with standard deviation (n=3).

**Table 3.**
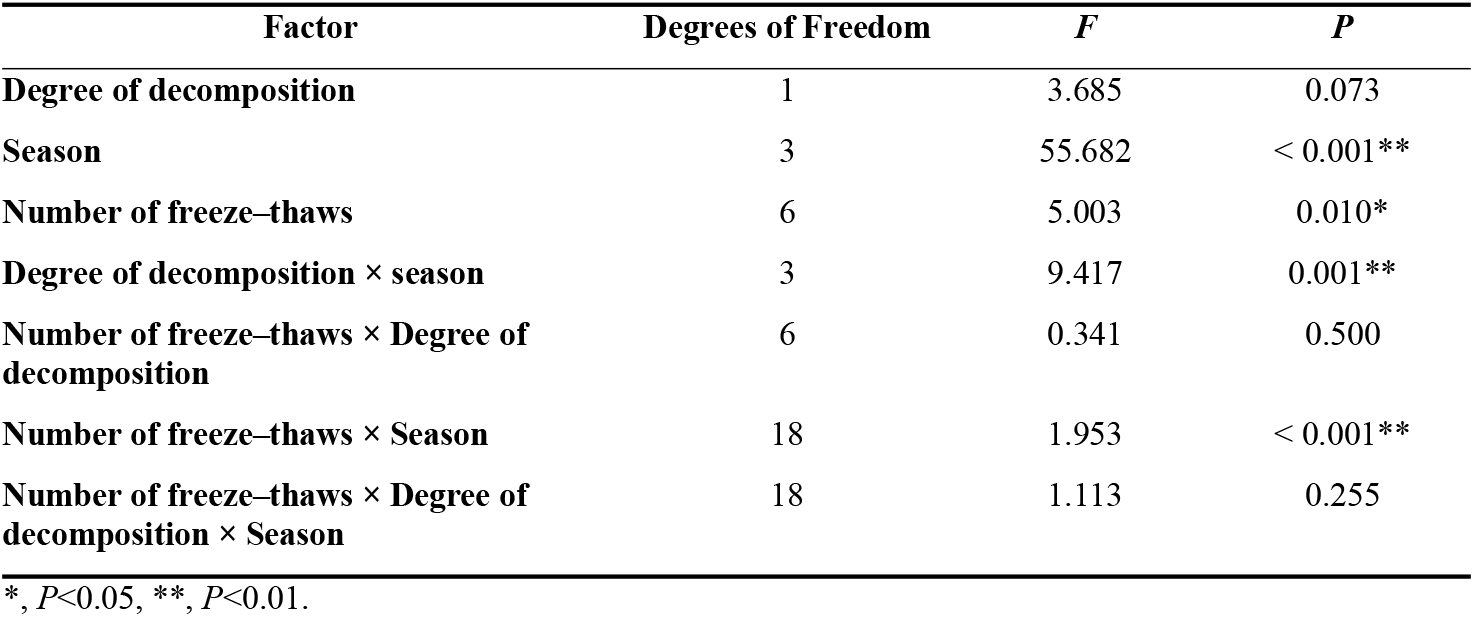
Seasonal variation in the organic carbon of dead litter at various levels of decomposition according to repeated-measures ANOVA.

| Factor | Degrees of Freedom | *F* | *P* |
| --- | --- | --- | --- |
| Degree of decomposition | 1 | 3.685 | 0.073 |
| Season | 3 | 55.682 | < 0.001** |
| Number of freeze–thaws | 6 | 5.003 | 0.010* |
| Degree of decomposition × season | 3 | 9.417 | 0.001** |
| Number of freeze–thaws × Degree of decomposition | 6 | 0.341 | 0.500 |
| Number of freeze–thaws × Season | 18 | 1.953 | < 0.001** |
| Number of freeze–thaws × Degree of decomposition × Season | 18 | 1.113 | 0.255 |

\*, $P < 0.05$ , \*\*, $P < 0.01$ .

Fig 3 shows that the number of freeze–thaws had a significant effect on litter organic carbon (P = 0.01). The interaction of freeze–thaw number and season had a significant effect on deadfall organic carbon (P<0.001). In the unfrozen season, the organic carbon of undecomposed litter after the 5th, 7th, and 30th freeze–thaws was significantly higher than the initial values, and the organic carbon of semi-decomposed litter was significantly higher for the 1st freeze–thaw than the 0th, 3rd, and 5th freeze–thaws. Organic carbon was significantly higher after the 30th freeze–thaw than after the 3rd freeze–thaw. In the freeze–thaw season, the organic carbon of undecomposed deadfall was significantly lower after the 1st and 7th freeze–thaws than initial values; the organic carbon of semi-decomposed deadfall was also higher after the 30th freeze–thaw than after the 1st and 5th freeze–thaws. The frequency of freeze–thaw cycles during the frozen season had no effect on the organic carbon content of deadfall. The organic carbon of semi-decomposed litter was significantly lower after the 5th freeze–thaw in the thaw season compared with the initial value, and the number of freeze– thaws had no effect on the organic carbon of undecomposed litter.

Organic carbon in the litter was significantly affected by season (P<0.001). The organic carbon of undecomposed litter was highest in the thaw season, followed by the frozen season and unfrozen season; the organic carbon of semi-decomposed litter was highest in the frozen season, followed by the thaw season and unfrozen season (Fig. 4). During the freeze–thaw season, the organic carbon of undecomposed litter decreased to a level below that of the frozen season; it then increased to a level above that of the frozen season but below that of the thaw season. By contrast, the organic carbon of semi-decomposed litter initially decreased to a level below that of the frozen season and the thaw season; it then increased to a level above that of the frozen season and thaw season. The mean organic carbon of undecomposed litter in the thaw season was 6.78% and 18% higher than that in the frozen and unfrozen seasons, respectively. The mean organic carbon of semi-decomposed litter in the thaw and unfrozen seasons was 6.34% and 28% lower, respectively, than that in the frozen season. The organic carbon of semi-decomposed litter in the freeze–thaw season was initially lower than that in the thaw season and the frozen season. the organic carbon of semi-decomposed litter increased from lower than 4.26% and 6.06% to greater than 6.83% and 1.93% in the thaw and frozen seasons, respectively.

**Fig 4.**
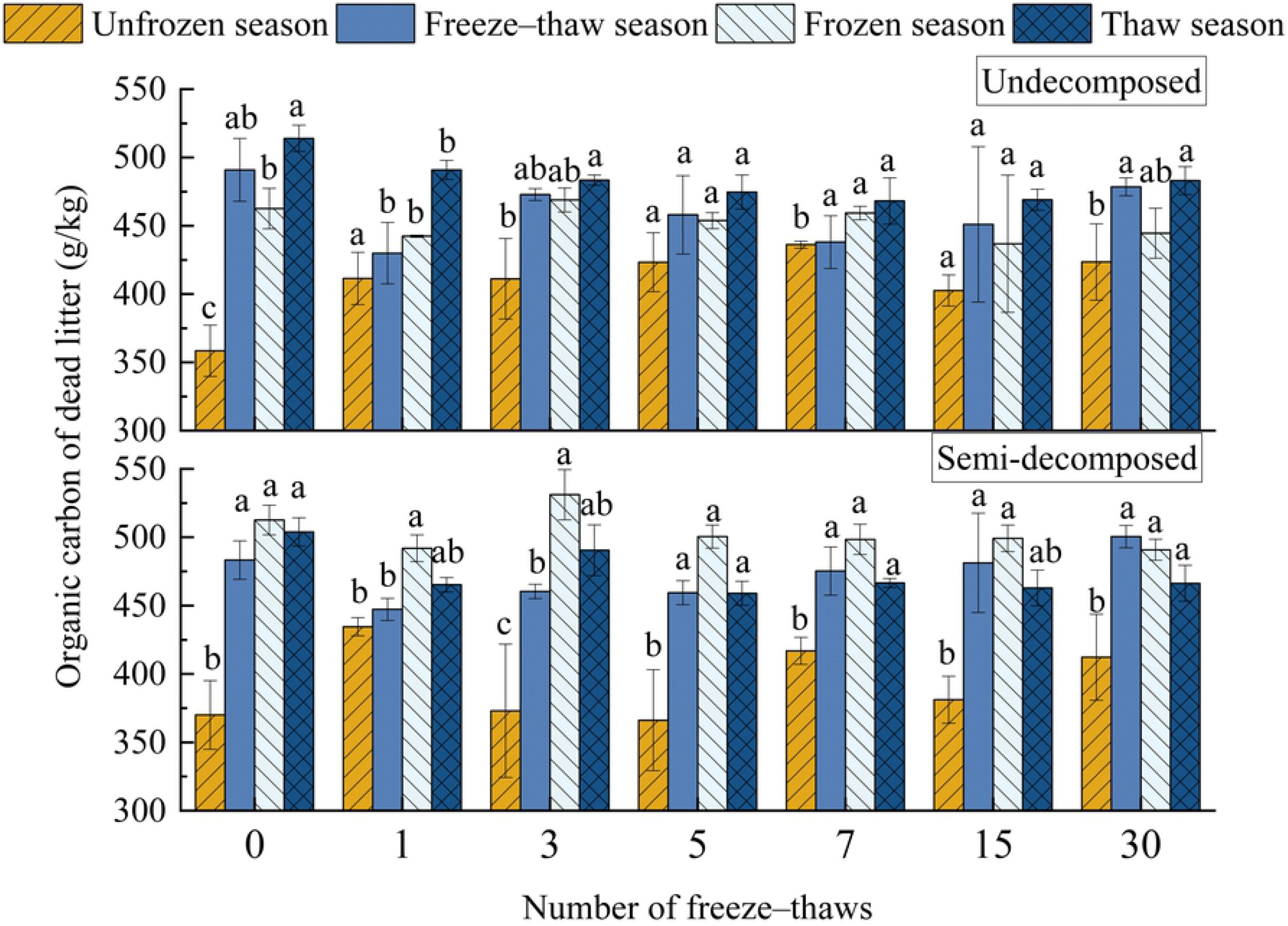
Effects of season on the carbon content of litter at various levels of decomposition under freeze–thaw treatment. Different lowercase letters indicate significant differences across seasons (*P*<0. 05); data in the graph are means with standard deviation (n=3).

### Effect of seasonal changes on the soil organic carbon of litter

The effect of season and number of freeze–thaws on soil organic carbon in deadfall was highly significant (P<0.01) (Table 4). The soil organic carbon of undecomposed litter was greater than the initial value during the freeze–thaw season after 30 freeze–thaws, and the opposite pattern was observed for semi-decomposed litter; the soil organic carbon of litter was lower in the freeze–thaw season compared with the other seasons. The soil organic carbon of the unfrozen and semi-decomposed layers of deadfall increased by 36.41% and 36.73%, respectively; the organic carbon in undecomposed deadfall soil decreased by 29.18%, and the organic carbon in semi-decomposed deadfall soil increased by 2.22%. During the frozen season, soil organic carbon in the undecomposed and semi-decomposed layers increased by 13.16% and 23.64%, respectively; during the thaw season, soil organic carbon in the undecomposed and semi-decomposed layers of deadfall increased by 9.68% and 25.35%, respectively.

**Table 4.** Seasonal variation in the soil organic carbon in the dead litter at various states of decomposition was studied using repeated-measures ANOVA.

| Factor | Degrees of freedom | *F* | *P* |
| --- | --- | --- | --- |
| Degree of decomposition | 1 | 0.133 | 0.720 |
| Season | 3 | 7.271 | 0.003** |
| Number of freeze-thaws | 6 | 7.319 | < 0.001** |
| Degree of decomposition × season | 3 | 0.096 | 0.961 |
| Number of freeze-thaws × Degree of decomposition | 6 | 1.432 | 0.210 |
| Number of freeze-thaws × Season | 18 | 3.271 | <0.001** |
| Number of freeze-thaws × Degree of decomposition × Season | 18 | 0.768 | 0.732 |

\*\* , $P < 0.01$ .

The interaction between the number of freeze–thaw cycles and season had a highly significant effect on soil organic carbon in the deadfall (P<0.001). Figure 5 shows that after the 7th and 30th freeze–thaw treatments during the unfrozen season, soil organic carbon in the undecomposed litter was significantly higher than the initial value; after the 7th, 15th, and 30th freeze–thaws, the soil organic carbon of semi-decomposed litter was considerably higher than that after the 0th and 5th freeze–thaw treatments. During the freeze–thaw season, soil organic carbon in the undecomposed litter was significantly lower after the 30th freeze–thaw treatment than after the 5th freeze–thaw treatment, and the number of freeze–thaw treatments had no effect on soil organic carbon in the semi-decomposed litter. During the frozen season, organic carbon in the deadfall was considerably higher after the 7th freeze–thaw than before the initial value and after the 5th freeze–thaw. The number of freeze–thaw cycles during the thaw season did not affect the soil organic carbon.

**Fig 5.**
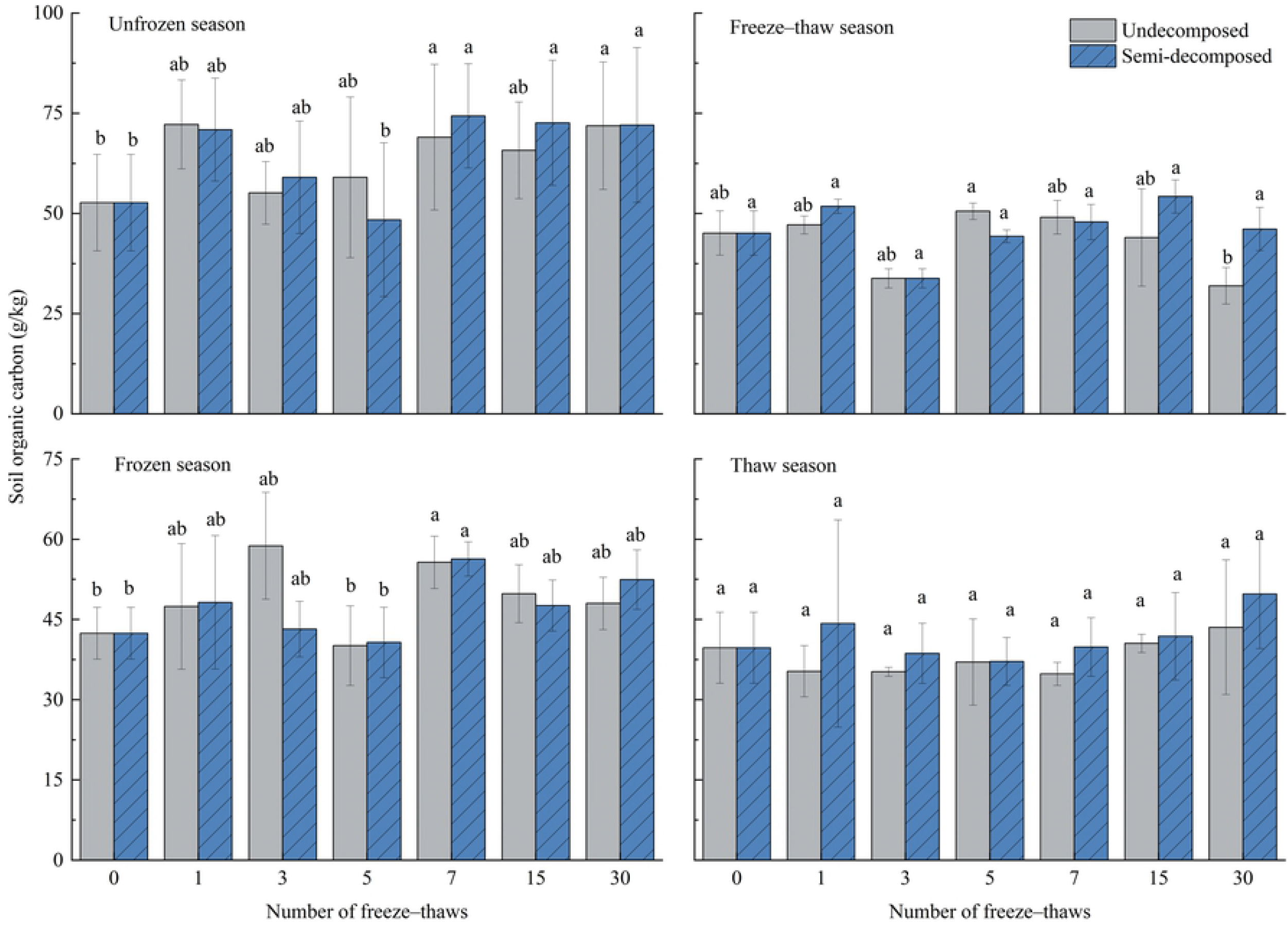
Effects of freeze–thaw time in different seasons on the soil carbon concentration at various degrees of litter decomposition. Significant differences in the number of freeze–thaws (*P*<0. 05) are indicated by different lowercase letters; data in the graph are means with standard deviation (n=3).

The effect of season on soil organic carbon was highly significant (P=0.003). Figure 6 shows that soil organic carbon in undecomposed deadfall soil was significantly higher in the 1st and 7th unfrozen seasons than in the thaw season, significantly higher in the 3rd frozen season than in the freeze–thaw and thaw seasons, and significantly higher in the 30th unfrozen season than in the freeze–thaw season. Soil organic carbon was significantly higher in semi-decomposed deadfall soil in the 3rd unfrozen season than in the freeze–thaw season, significantly higher in the 7th unfrozen season than in the freeze–thaw and thaw seasons, and significantly higher in the 15th unfrozen season than in the thaw season. The number of freeze–thaw seasons had no effect on the organic carbon of deadfall.

**Fig 6.**
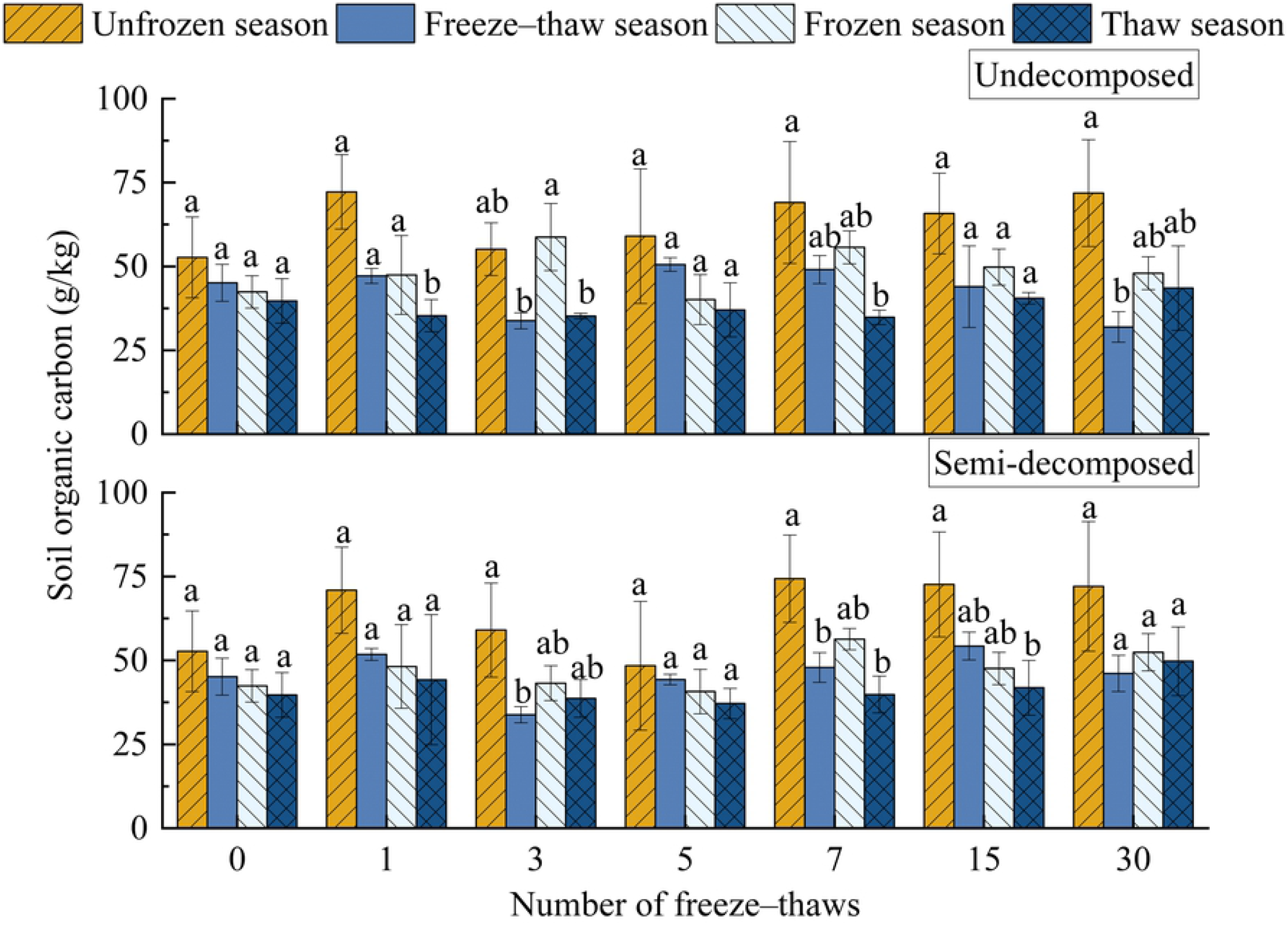
Effects of season on the carbon content of soil litter at various levels of decomposition under freeze–thaw treatment. Different lowercase letters indicate significant differences across seasons (*P*<0.05); data in the graph are means with standard deviation (n=3).

## Discussion

Forests are the most cost-effective carbon absorbers because they fix atmospheric CO2 in the form of organic matter in the plant body and soil through photosynthesis, thus reducing the accumulation of greenhouse gases. Forests thus play roles in the global carbon cycle as a source, reservoir, and sink, and any increase or decrease in their carbon stocks can affect variation in atmospheric CO2 concentrations [26]. All organic matter created by biological components of a forest ecosystem and returned to the forest floor as a source of material and energy for decomposers to sustain ecosystem function is referred to as apoplankton. The biogeochemical cycle of forest ecosystems includes the breakdown of apoplastic matter, and the rate of decomposition of this matter has a substantial effect on the productivity of ecosystems. Deadfall decomposition also has a considerable effect on the physicochemical quality of soil and is a primary predictor of the biomass and nutrient content of the forest floor [27]. Forest litter is an important component of the biomass and material cycle of forest ecosystems, as it releases nutrients and energy fixed by forest plants from the soil and atmosphere to the environment through microbial decomposition. It is thus a major link in the energy flow and material cycle of forest ecosystems and one of the most important media for carbon exchange between the atmosphere and soil [28]. The forest floor is subjected to long periods of seasonal freezing and thawing at high latitudes and altitudes, which affects the breakdown of apoplastic material by mechanical fragmentation, the leaching of the material’s physical structure, as well as the leaching of biological constituents [29,30]. The frequency of freeze–thaw cycles has a substantial effect on the composition of deadfall. The number of freeze–thaw cycles had a substantial effect on the mass loss of litter (P<0.05). The rate of the mass loss of litter was high from the start of the 1st to the 7th freeze–thaw in the unfrozen season (Fig. 1 and 2), indicating that the residual rate was much lower after the 7th freeze–thaw than the initial value. The rate of mass loss of semi-decomposing litter decreased, increased, and then decreased, and the rate of mass loss peaked after the 7th freeze–thaw; the residual rate of semi-decomposing litter was significantly lower after the 7th freeze–thaw than its initial value. Because of the higher temperatures and greater mass-loss rates in the unfrozen season, the overall rate of mass loss was higher in the unfrozen season than in the other three seasons. Because the temperature fluctuates above and below 0°C during the freeze–thaw season, the overall rate of mass loss of undecomposed litter was higher than that of semi-decomposed litter. The physical fragmentation of undecomposed litter during the freeze–thaw season enhanced the rate of mass loss of undecomposed litter. The effect of freeze–thaws on the semi-decomposed litter was minimal. The rate of mass loss was not lower in the frozen season than in other seasons. In the frozen season, the residual rate was much lower after the 5th and 15th freeze–thaw cycles than its initial value. This suggests that mass loss still occurs during the frozen season. The lowest rate of mass loss was observed during the thaw season. The number of freeze–thaw treatments did not affect the litter mass retention rate throughout the thaw season. The litter present in the thaw season did not decompose in both the freeze–thaw and frozen seasons; in other words, the litter present in the thaw season was litter that does not easily decompose. Alternatively, the water in the litter was damaged by the low temperature during the frozen season. Fluctuations in temperature above and below 0°C throughout the thaw season are insufficient for disintegrating the leaves. This is consistent with the results of Gao J [31]. showing that the rate of decomposition of broadleaf forest litter is highest in the early phases of decomposition and gradually decreases thereafter.

Apoplankton is a vital component of ecosystems, and characterizing the volume of apoplankton and its effect on nutrient dynamics is important for understanding the material cycles and energy flows of ecosystems [32,33]. Some of the carbon in decomposing apoplankton is directly released into the atmosphere, whereas the rest enters the soil, becomes part of the soil carbon cycle, or is released into the atmosphere via soil respiration [28]. The pace of apoplastic matter decomposition and conversion reflects the rate of nutrient return and cycling on the forest floor [11]. The release of nutrients from decaying litter is primarily controlled by biotic and abiotic processes and includes leaching-enrichment-release, enrichment-release, and direct release [34]. Increasing temperatures have a substantial impact on the structure, function, dynamics, and distribution of forest vegetation, and these changes can affect the carbon sequestration capacity of forest vegetation [35–38]. The significance of temperature regulation is intimately linked to the carbon cycle processes associated with the buildup and breakdown of apoplankton [2]. Climate change has direct and indirect effects on the decomposition process [39,40] and is the most important ecological element controlling apoplankton decomposition [41]. Carbon and nutrient cycling in cold locations is affected by the mass loss and nutrient release of forest litter during the freeze–thaw season [42]. Season, the number of freeze–thaw cycles, as well as their interaction had significant effects on the organic carbon in litter (P<0.01). The organic carbon of deadfall increased at the end of the culture period in the unfrozen season and the frozen and thaw seasons; the organic carbon of the undecomposed dead litter decreased at the end of the culture period in the freeze–thaw season; and the organic carbon of the semi-decomposed dead litter increased at the end of the culture period in the freeze–thaw season. Organic carbon was higher later in the freeze–thaw stage of the unfrozen season compared with earlier in that season. After the 7th freeze–thaw in the freeze–thaw season, the organic carbon of undecomposed deadfall began to increase; the organic carbon of semi-decomposed deadfall gradually increased after the 1st freeze–thaw. The organic carbon of semi-decomposed litter in the 30th freeze–thaw cycle was substantially higher than that in the 1st freeze-thaw cycle; the number of freeze–thaws had no effect on the organic carbon of deadfall in the frozen season. The organic carbon of semi-decomposed deadfall was significantly lower than the initial value after the 5th freeze–thaw in the thaw season, which might stem from the state of decomposition of the deadfall and the presence of several carbon elements that had not yet been released into the soil. The number of freeze–thaws had no effect on the organic carbon of undecomposed deadfall in the thaw season. The organic carbon of the litter was significantly affected by the interaction between the degree of litter decomposition and season (P=0.001). The organic carbon of undecomposed dead litter was highest in the thaw season, followed by the freeze–thaw season, frozen season, and unfrozen season, and the organic carbon of semi-decomposed dead litter was highest in the frozen season, followed by the thaw season, freeze–thaw season, and unfrozen season. The organic carbon of the litter began to fix carbon at the beginning of the unfrozen season, and organic carbon increased in the freeze–thaw season. After the freeze– thaw season, the organic carbon of the undecomposed litter decreased, and the organic carbon of the semi-decomposed litter increased. The carbon of the undecomposed litter was transferred to the semi-decomposed litter, and this caused the organic carbon of the undecomposed litter to decrease in the frozen season. The soil organic carbon began to rise during the thaw season, given that the carbon of semi-decomposed litter carbon was released into the soil. In conclusion, the carbon stock of dead litter was lowest during the unfrozen season. This is consistent with the results of Yuan W.Y [43] showing that litter carbon stocks are lower in the summer, as well as those of Zhou Y.B [20] showing that freeze–thaws promote the accumulation of organic carbon in the residual litter.

In terrestrial ecosystems, the soil is the largest carbon pool, but its turnover is slow. The organic and inorganic carbon pools, as well as a small part of the soil inorganic carbon pool, comprise this system [44]. Soil organic carbon is an important component of the carbon pools of terrestrial ecosystems, and slight changes in soil organic carbon can have substantial effects on atmospheric greenhouse gas concentrations [45]. Approximately 1400–1500 Gt of carbon is stored in organic form in soils around the world, which is two to three times greater than the carbon pool in terrestrial vegetation (500–600 Gt) and more than twice as much as the carbon pool in the atmosphere (750 Gt) [46]. Forest soil contains approximately 40% of the global terrestrial soil organic carbon [47]. The amount of forest soil carbon is determined by the relationship between the amount of biomass input, the amount of carbon released by decomposition, and the amount of carbon lost into the water system; it is derived from the input of aboveground and belowground apoplastic material and the decomposition of organic matter and is in the form of decomposed plant and animal remains, debris, or organic matter [48]. As a result, the size of the soil carbon pool is affected not only by the vegetation but also by local climatic conditions [49]. Climate change is linked to the decomposition and buildup of organic carbon in forest soils [50]. The accumulation of soil organic carbon is affected by various factors, including climate. Climatic conditions determine the type of vegetation, its productivity, and thus the amount of organic carbon input to the soil. Microorganisms are the primary drivers of soil organic carbon decomposition and turnover, and climate can affect the decomposition and transformation of organic carbon by microorganisms by altering soil moisture, soil aeration, and temperature [51–54]. Freeze–thaws are a common natural phenomenon in which soil freezes and thaws as a result of variation in temperature [55]. They are particularly common in the northern hemisphere, where temperatures fluctuate around 0°C; these freezing and thawing conditions affect the physical, chemical, and biological properties of the soil. In Northeast China, freeze–thaw cycles occur for approximately six months a year and thus have a substantial effect on soil organic carbon turnover [56,57]. The release of organic carbon from the soil is also accelerated by the melting of permafrost due to global warming. The depth and duration of the thaw determine the stability of the soil organic carbon pool during freeze–thaw conditions [58]. The physical breakdown of apoplankton increases with the number of freeze– thaw cycles, which promotes apoplankton decomposition. Apoplankton is an important component of the forest ecosystem carbon pool, and its decomposition plays a critical role in the formation of soil organic matter and the forest carbon cycle [2,59]. The number of seasons and freeze–thaw cycles, as well as their interaction, had a substantial effect on the soil organic carbon content in litter at different levels of decomposition (P<0.01). Soil organic carbon accumulates after the end of the unfrozen season due to photosynthesis, the high carbon sequestration ability of plants, and carbon fixation in the soil, and organic carbon was higher at this point compared with the initial value in the freeze–thaw phase (after the 7th freeze–thaw). During the freeze– thaw season, the organic carbon of undecomposed litter soil decreased, and the organic carbon of semi-decomposed litter slightly increased; the organic carbon of undecomposed litter soil was the highest in the 5th freeze–thaw season (50.6±2.0 g/kg), and the organic carbon of undecomposed litter soil was significantly lower after 30 freeze–thaws (31.9±4.6 g/kg) than after the 5th freeze–thaw. This is likely explained by the release of the organic carbon of undecomposed litter into the soil through the decomposition of litter; furthermore, the carbon absorbed by the soil from semi-decomposed litter was less than the carbon released into the atmosphere by the soil. Soil organic carbon increased in both the frozen and thaw seasons, and it was higher after the 7th compared with the initial value and the value after the 5th frozen season. This might be explained by the slow release of organic carbon from deadfall into the soil. This result is consistent with the organic carbon transfer process of deadfall described above, which reflects the migration of carbon between deadfall and soil over time. Figure 6 shows that soil organic carbon levels were often higher in the unfrozen season than in the other three seasons. Soil organic carbon was considerably higher in undecomposed deadfall soils in the 1st, 3rd, and 7th unfrozen seasons than in the thaw season; it was also significantly higher in the 3rd and 30th unfrozen seasons than in the freeze–thaw season. The soil organic carbon of semi-decomposed soil was significantly higher in the 3rd and 7th unfrozen seasons than in the freeze–thaw season; it was also significantly higher in the 7th and 15th unfrozen seasons than in the thaw season. Soil organic carbon was highest in the unfrozen season, freeze–thaw season, frozen season, thaw season to unfrozen season, frozen season, thaw season, and freeze–thaw season after 30 freeze–thaw cycles. This might stem from the fact that plants have a greater potential to sequester carbon during the unfrozen season, which is consistent with the lower organic carbon in the unfrozen season compared with the other three seasons. The progressive degradation of undecomposed litter into semi-decomposed litter, where organic carbon is retained in the litter layer and little carbon is released into the soil, might explain the decrease in soil organic carbon during the freeze–thaw season. The increase in soil organic carbon during the frozen season most likely stems from the increase in organic carbon in the litter and lower CO2 evaporation, which results in a faster rate of carbon migration into the soil. Because the initial organic carbon value is lower and more carbon is trapped in the litter, the organic carbon was often lower in the thaw season than in the other three seasons. Chen Z.H [60] found that organic carbon levels decrease from January to March and are significantly higher in subsequent seasons.

## Conclusions

The temperature was higher, the mass-loss rate was faster, and the overall mass-loss rate of litter was higher in the unfrozen season than in the other three seasons. This resulted in greater litter organic carbon and soil organic carbon in the unfrozen season because of the strong carbon sequestration capacity of plants. The temperature fluctuates above and below 0°C during the freeze–thaw season, which physically breaks the undecomposed litter and speeds up its mass-loss rate. This results in increases in the organic carbon of undecomposed litter and decreases in the soil organic carbon of undecomposed litter; the opposite patterns were observed for the organic carbon of semi-decomposed litter and soil. Some mass loss of litter was observed during the frozen season. The mass-loss rate of litter during the thaw season was the lowest; litter organic carbon decreased and soil organic carbon increased in both seasons. The organic carbon of undecomposed litter was highest in the thaw season, followed by the freeze–thaw season, frozen season, and non-frozen season, the organic carbon of semi-decomposed litter was highest in the frozen season, followed by the thaw season, freeze–thaw season, unfrozen season to freeze–thaw season, frozen season, thaw season, and unfrozen season. After freeze–thaw treatment, soil organic carbon was highest in the unfrozen season, freeze–thaw season, frozen season, thaw season to unfrozen season, frozen season, thaw season, and freeze– thaw season.

## Acknowledgments

The authors thank the forestry workers of the Dailing Forestry Bureau for their help in establishing the plots. We also thank the teachers and students of the Department of Forest Engineering, Northeast Forestry University (NEFU), China for providing and collecting the data for this study.

## References

1. Wang Y, Liu JS, Wang GP, Zhou WM. Study on the Effect of Freezing and Thawing Action to Soil Physical and Chemical Characteristics. Geography and Geo-Information Science. 2007;23(2):91–6.

2. Yang WQ, Deng RJ, Zhang J. Forest litter decomposition and its responses to global climate change. Chinese Journal of Applied Ecology. 2007;18(12):2889–95.

3. Wei J, Wu G, Deng H. Researches on nutrient return of litterfall in the alpine tundra ecosystem of Changbai Mountains. Acta Ecologica Sinica. 2004;24(10):2211–6. PubMed PMID: CSCD:1645850.

4. Jing G, Liu B, Rong J, Sun X. Distribution and Characteristics of Freeze-thaw Erosion in Heilongjiang Province. Bulletin of Soil and Water Conservation. 2016;36(4):320–5. PubMed PMID: CSCD:5784474.

5. Zhongmin W, Yide L, Qingbo Z, Guangyi Z, Bufeng C, Zhihu D, et al. Carbon pool of tropical mountain rain forests in Jianfengling and effect of clear-cutting on it. Chinese Journal of Applied Ecology. 1998;9(4):341–4.

6. Shan XS, Yao SB. Analysis on Regional Differences of Forest Carbon Storage in China Based on Biomass Expansion Factor. Journal of Beijing Forestry University(Social Sciences). 2009;8(03):109–14.

7. Post WM, Emanuel WR, Zinke PJ, Stangenberger AG. Soil carbon pools and world life zones. Nature. 1982;298(5870):156–9. doi: 10.1038/298156a0.

8. Kramer PJ. Carbon Dioxide Concentration, Photosynthesis, and Dry Matter Production. Bioscience. 1981;(1):29–33.

9. Scott NA, Binkley D. Foliage litter quality and annual net N mineralization: comparison across North American forest sites. Oecologia. 1997;111(2):151–9.

10. Wu CZ, Hong W, Jiang ZL, Zheng FH. Advances in Research of Forest Litter-fall in China. Acta Agriculturae Universitatis JiangxiensisNatural Science Edition. 2000;22(3):405. PubMed PMID: CSCD:681087.

11. Fangli SU, Mingguo LIU, Dexia CHI, Xiangwen K, Wanliang HU. Effect of Different Thinning Intensity on the Properties of Litter. Journal of Soil Science. 2007;38(6):1096–9. PubMed PMID: CSCD:3041546.

12. Wu F, Yang W, Zhang J, Deng R. Litter decomposition in two subalpine forests during the freeze–thaw season. Acta Oecologica. 2010;36(1):135–40. doi: 10.1016/j.actao.2009.11.002.

13. Wen L, Lei P, Xiang W, Yan W, Liu S. Soil microbial biomass carbon and nitrogen in pure and mixed stands of Pinus massoniana and Cinnamomum camphora differing in stand age. Forest Ecology & Management. 2014;328:150–8.

14. Grogan P, Michelsen A, Ambus P, Jonasson S. Freeze-thaw regime effects on carbon and nitrogen dynamics in sub-arctic heath tundra mesocosms. Soil Biology & Biochemistry. 2004;36(4):641–54.

15. Jia G, Zhou Y, Dai L, Zhou W. Effect of freezing-thawing on the carbon and nitrogen mineralization in Changbai Mountain. Ecology and Environmental Sciences. 2012;21(4):624–8. PubMed PMID: CSCD:4553509.

16. Jenkinson DS, Adams DE, Wild A. Model estimates of CO2 emissions from soil in response to global warming. Nature. 1991;351(6324):304–6.

17. Li Q, Zhou D, Chen X. The accumulation,decomposition and ecological effects of above-ground litter in terrestrial ecosystem. Acta Ecologica Sinica. 2014;34(14):3807–19. PubMed PMID: CSCD:5202278.

18. Chen T, Xi M, Kong F, Li Y, Pang L. A review on litter decomposition and influence factors. Chinese Journal of Ecology. 2016;35(7):1927–35. PubMed PMID: CSCD:5744896.

19. Devi NB, Yadava PS. Influence of climate and litter quality on litter decomposition and nutrient release in sub-tropical forest of Northeast India. Journal of Forestry Research. 2010;21(2):143–50.

20. Zhou YB, Jia GJ, Zhou WM, Guo YT. Study on Decomposition Dynamics of Carbon of Changbai Mountain Litter. Northern Horticulture. 2012;(11):32–4.

21. Zhou R, Zhang Y, Peng M, Jin Y, Song Q. Effects of Climate Change on the Carbon Sequestration Potential of Forest Vegetation in Yunnan Province, Southwest China. Forests. 2022;13(2):306. PubMed PMID: doi:10.3390/f13020306.

22. Chiang JM, Iverson LR, Prasad A, Brown KJ. Effects of Climate Change and Shifts in Forest Composition on Forest Net Primary Production. J Integr Plant Biol. 2008;50(11):1426–39. doi: 10.1111/j.1744-7909.2008.00749.x. PubMed PMID: WOS:000260336800010.

23. Fei XH, Song QH, Zhang YP, Liu YT, Sha LQ, Yu GR, et al. Carbon exchanges and their responses to temperature and precipitation in forest ecosystems in Yunnan, Southwest China. Sci Total Environ. 2018;616:824–40. doi: 10.1016/j.scitotenv.2017.10.239. PubMed PMID: WOS:000424121800084.

24. Zhang Y, Hu H. Study on Effects of Forest Litter on Soil Physicochemical Properties and Carbon Library. Journal of Green Science and Technology. 2018;(10):9–10+3. doi: 10.16663/j.cnki.lskj.2018.10.004.

25. Sun K, Li J, Yang G, Zuo M, Hu J. Effect of alterations in forest litter inputs on soil C and N storage distribution in Pinus yunnanensis forest in central Yunnan Plateau. Acta Ecologica Sinica. 2021;41(8):3100–10. PubMed PMID: CSCD:6956974.

26. Hopmans JW. The Potential of U.S. Forest Soils to Sequester Carbon and Mitigate the Greenhouse Effect. Vadose Zone Journal. 2004;3(2):734–5. doi: https://doi.org/10.2136/vzi2004.0734.

27. Wang J, Huang, Jianhui. Comparison of major nutrient release patterns in leaf litter decomposition in warm temperate zone of china. Acta Phytoecological Sinica. 2001;25(3):375–80. PubMed PMID: 540919.

28. Lei PF. A preliminary study on carbon storage in soil and its formation mechanism in Chinese fir plantation [Master]: Central South Forestry College; 2004.

29. Sulkava P, Huhta V. Effects of hard frost and freeze-thaw cycles on decomposer communities and N mineralisation in boreal forest soil. Applied Soil Ecology. 2003;22(3):225–39.

30. Withington CL, Sanford RL. Decomposition rates of buried substrates increase with altitude in the forest-alpine tundra ecotone. Soil Biology & Biochemistry. 2007;39(1):68–75. doi: 10.1016/j.soilbio.2006.06.011. PubMed PMID: WOS:000242720300006.

31. Gao J, Wei X, Chen Y, Dong Y, Yang Y, Zhang D. Litter decomposition rates and organic carbon dynamics in subalpine forest during freeze-thaw cycles. Acta Ecologica Sinica. 2021;41(9):3734–43. PubMed PMID: CSCD:6966162.

32. Witkamp M. Microbial Populations of Leaf Litter in Relation to Environmental Conditions and Decomposition. Ecology. 1963;44(2).

33. Maguire DA. Branch mortality and potential litterfall from Douglas-fir trees in stands of varying density. Forest Ecology and Management. 1994;70(1):41–53. doi: https://doi.org/10.1016/0378-1127(94)90073-6.

34. Zhang D. Advances of Enzyme Activities in the Process of Litter Decomposition. entia Silvae Sinicae. 2006;42(01):105–9.

35. Guo P, Zhao X, Shi J, Huang J, Zeng J. The influence of temperature and precipitation on the vegetation dynamics of the tropical island of Hainan. Theoretical and Applied Climatology. 2021;143(1-2):1–17.

36. Watson RT. Technical summary: Impacts, adaptations and mitigation options. Climate Change. 1996.

37. Hui D, Deng Q, Tian H, Luo Y. Climate Change and Carbon Sequestration in Forest Ecosystems. In: Chen W-Y, Suzuki T, Lackner M, editors. Handbook of Climate Change Mitigation and Adaptation. New York, NY: Springer New York; 2016. p. 1–40.

38. Weng ES, Zhou GS. Modeling distribution changes of vegetation in China under future climate change. Environ Model Assess. 2006;11(1):45–58. doi: 10.1007/s10666-005-9019-1. PubMed PMID: WOS:000234555400004.

39. Park S, Cho KH. Nutrient leaching from leaf litter of emergent macrophyte (Zizania latifolia) and the effects of water temperature on the leaching process. Korean Journal of Biological Sciences. 2003;7(4):289–94.

40. DeLucia, Hamilton, Naidu, Thomas, Andrews, Finzi, et al. Net primary production of a forest ecosystem with experimental CO2 enrichment. Science (New York, NY). 1999;284(5417):1177–9. doi: 10.1126/science.284.5417.1177. PubMed PMID: MEDLINE:10325230.

41. Vitousek PM, Turner DR, Parton WJ, Sanford RL. Litter Decomposition on the Mauna Loa Environmental Matrix, Hawai’i: Patterns, Mechanisms, and Models. Ecology. 1994;75(2):418–29. doi: https://doi.org/10.2307/1939545.

42. Wu F, Yang W, Zhang J, Deng R. Litter decomposition in two subalpine forests during the freeze–thaw season. Canadian Journal of Forest Research. 2010;36(1):135–40.

43. Yuan WY, Xian-Wei U, Zhang J, Rong L. Preliminary Studies on Carbon Reserves of Litterfall and Fine Root in an Age Series of Eucalyptus grandis Plantation. Forest Research. 2009;22(3):385–9.

44. Schlesinger W. Carbon storage in the Caliche of arid soils: A case study from Arizona. Soil Science – SOIL SCI. 1982;133:247–55. doi: 10.1097/00010694-198204000-00008.

45. Kimble JM, Lal R, Follett R. Agricultural Practices and Policies for Carbon Sequestration in Soil 2016. 1–512 p.

46. Schlesinger WH. Evidence from chronosequence studies for a low carbon-storage potential of soils. Nature. 1990;348(6298):232–4. doi: 10.1038/348232a0.

47. Angelika T, Nina B, Ernst-Detlef S. Carbon stocks and soil respiration rates during deforestation, grassland use and subsequent Norway spruce afforestation in the Southern Alps, Italy. Tree Physiology. 2000;(13):849–57.

48. Effect of Forest on Atmosphere Carbon Balance. WORLD FORESTRY RESEARCH. 1997.

49. Wang SQ, Zhou CH, Liu J, Li KR, Yang XM. Simulation analyses of Terrestial Carbon Cycle Balance model in Northest China. Acta Geographica Sinica. 2001;56:390–400.

50. Zhou X-y, Zhang C-y, Guo G-f. [Effects of climate change on forest soil organic carbon storage: a review]. Ying Yong Sheng Tai Xue Bao. 2010;21(7):1867–74. PubMed PMID: 20879549.

51. Fitzhugh RD, Driscoll CT, Groffman PM, Tierney GL, Fahey TJ, Hardy JP. Effects of soil freezing disturbance on soil solution nitrogen, phosphorus, and carbon chemistry in a northern hardwood ecosystem. Biogeochemistry. 2001;56(2):215–38.

52. Herrmann A, Witter E. Sources of C and N contributing to the flush in mineralization upon freeze-thaw cycles in soils. Soil Biology & Biochemistry. 2002;34(10):1495–505.

53. Sharma S, Szele Z, Schilling R, Munch JC, Schloter M. Influence of Freeze-Thaw Stress on the Structure and Function of Microbial Communities and Denitrifying Populations in Soil. Applied and Environmental Microbiology. 2006;72(3):2148–54.

54. Davidson EA, Trumbore SE, Amundson R. Soil warming and organic carbon content. Nature. 2000;408(6814):789–90. doi: 10.1038/35048672.

55. Piao H, Liu G, Hong Y. Effects of alternative drying-rewetting and freezing-thawing on soil fertility and ecological environment. Chinese Journal of Ecology. 1995;14(6):29–34. PubMed PMID: 284603.

56. Wang Z, Zhang YL, Zhang L, Fan QF, Liu C. Effect of Freeze/Thawing Frequency and Soil Water Content on Total Organic Carbon and Dissolved Organic Carbon of Burozem. Journal of Agro-Environment Science. 2012.

57. Chang Z, Ma Y, Liu W, Feng Q, Su Y, Xi H, et al. Effect of soil freezing and thawing on the carbon and nitrogen in forest soil in the Qilian Mountains. Journal of Glaciology and Geocryology. 2014;36(1):200–6. PubMed PMID: CSCD:5103479.

58. Goulden, Wofsy, Harden, Trumbore, Crill, Gower, et al. Sensitivity of boreal forest carbon balance to soil thaw. Science (New York, NY). 1998;279(5348):214–7. doi: 10.1126/science.279.5348.214. PubMed PMID: MEDLINE:9422691.

59. Chen Z, Shen Y, Tan B, Li H, You C, Xu Z, et al. Decreased Soil Organic Carbon under Litter Input in Three Subalpine Forests. Forests. 2021;12(11):1479. PubMed PMID: doi:10.3390/f12111479.

60. Chen Z, Jiao Z, Liu Y, Xu Z, Tan B, Zhang L. Influences of seasonal litter input on soil active organic carbon in subalpine forests in western Sichuan. Chinese Journal of Applied and Environmental Biology. 2021;27(3):594–600. PubMed PMID: CSCD:7004492.

